# MEK1/2 kinases cooperate with c-Myc:MAX to prevent polycomb repression of *TERT* in human pluripotent stem cells

**DOI:** 10.1101/2024.09.16.613267

**Authors:** Spandana Kotian, Lindsay F. Rizzardi, Josh L. Stern

## Abstract

Telomerase counteracts telomere shortening, enabling human embryonic stem cells (hESC) to undergo long-term proliferation. MAPK signaling plays a major role in regulating the self-renewal of hESC, and previous studies in induced pluripotent stem cells (iPSC) suggested that expression of *TERT*, the gene encoding the catalytic subunit of telomerase, relies on MAPK signaling. We examined whether MEK-ERK signaling regulated *TERT* transcription in a model of normal hESC. Kinase inhibitors of MEK1 and MEK2 (MEKi) or ERK1 and ERK2 (ERKi) significantly repressed *TERT* mRNA levels. Using chromatin immunoprecipitation (ChIP) we observed that MEKi induced the accumulation of the repressive histone mark histone 3 lysine 27 trimethylation (H3K27me3) at the *TERT* proximal promoter. This increase corresponded with a loss of histone 3 lysine 27 acetylation (H3K27ac) which is associated with transcriptionally active loci. Inhibition of the polycomb repressive complex 2 (PRC2), which deposits H3K27me3, partially rescued the loss of *TERT* expression, indicating that MEK1/2 activity can limit PRC2 activity at *TERT*. Inhibition of MEK/ERK kinases also repressed expression of c-Myc, a transcription factor reported to regulate *TERT* in other immortalized cells. Consistent with a key role for c-Myc in regulating *TERT*, low doses of a c-Myc:MAX dimerization inhibitor induced a striking and rapid gain of H3K27me3 at *TERT* and repressed *TERT* transcription in hESC. Inhibiting c-Myc:MAX dimerization also resulted in lower MAX recruitment to *TERT*, suggesting that this complex acts in cis at *TERT*. Our study using a model of normal human pluripotent stem cells identifies new regulators and mechanisms controlling transcription of an important, developmentally regulated gene involved in telomere protection.

## INTRODUCTION

Telomeres are protective protein-DNA structures at chromosome termini. Telomerase is an enzyme that recruits to telomeres to counteract chromosome shortening caused by DNA replication problems at the 3’ end [1–3]. The catalytic subunit of the telomerase complex, encoded by *TERT*, is a reverse transcriptase that adds hexameric DNA repeats to telomeric ends [4,5]. During development, telomerase activation occurs at the blastocyst stage of embryogenesis, following which the expression of *TERT* becomes restricted to stem cell compartments [6]. While the mature telomerase enzyme complex contains several factors, components besides *TERT* are expressed ubiquitously [7,8]. Thus, regulation of *TERT* transcripts is the primary on/off switch for telomerase activity. There are significant gaps in our knowledge of the regulation *TERT* transcripts in stem cells and, therefore, the establishment of telomere length during embryogenesis and the maintenance of their long-term proliferative capacity.

Haploinsufficiency of *TERT* and other factors involved in telomere maintenance cause telomere biology disorders that can present as segmental progeriatric diseases such as dyskeratosis congenita, idiopathic pulmonary fibrosis and cardiomyopathy in Duchenne muscular dystrophy [9]. In addition, stem cell function in aging tissues is impaired by telomere shortening and studies suggest that early onset age-associated morbidities might be addressed by improving telomere maintenance (reviewed in [10]).

Despite the critical role of *TERT* in human embryogenesis and normal cellular aging, there are few studies of the regulation of *TERT* transcripts in normal human stem cells. Although mouse and human *TERT* promoters share partial sequence homology, mouse *TERT* undergoes different developmental regulation compared to human telomerase [11]. Indeed, most murine adult tissues express telomerase [12]. Because of this, studies of transcriptional regulation of *TERT* in murine models have a limited capacity to capture the dynamics of human *TERT* developmental regulation. *TERT* expression in human pluripotent stem cells is controlled by cell density and alternative splicing which developmentally regulates *TERT* mRNA stability [13,14]. The human *TERT* promoter contains potential binding sites for several transcription factors including pluripotency-related factors such as c-Myc and KLF4. KLF4 has been shown to bind to and activate the *TERT* proximal promoter in human stem cells [15].

One limitation to studying *TERT* transcriptional regulation in pluripotent cells has been the slower development of human stem cell culture models. An early model of human pluripotent cells is the induced pluripotent stem cell (iPSC). An essential feature of reprogramming somatic cells, which lack telomerase expression, into iPSC involves the reactivation of *TERT* transcription and the restoration of telomere length [16,17]. However, it remains unknown if these artificial stem cells accurately model the regulation of *TERT* in normal stem cell. In recent years, several advanced cell culture models have been developed to recapitulate the naïve and primed states of human embryonic development [18–20] providing the first opportunity to study *TERT* regulation in models of normal human pluripotent stem cells.

In contrast to murine embryonic stem (mES) cells where partial inhibition of MEK1/2 kinase activity is essential for maintaining self-renewal capacity [21], human embryonic stem cells (hESC) rely on MEK-ERK signaling to promote self-renewal and pluripotency [22]. In this study, we show in a model of normal human pluripotent stem cells that *TERT* transcription is regulated through MEK/ERK signaling and Myc:MAX heterodimers. The implications of this study are that human stem cells are likely to rely on these factors to maintain telomere length and their long-term proliferative capacity.

## 2. MATERIALS AND METHODS

### Cell culture

Human pluripotent embryonic stem cells (WA01) were obtained from Wicell Inc. and induced pluripotent stem cells (iPSC) derived from reprogramming CD34^+^ hematopoietic cells (BXS0116) were obtained from American Type Cell Culture Collection. These cells were maintained in feeder-free culture on vitronectin (Gibco, A14700) in Essential 8 Flex media (Gibco, A2858501), supplemented with glutamine analog SG-200 (Cytiva, SH30590.01) and 1% Penicillin-Streptomycin (Cytiva, SV30010). For passaging, cells were dissociated using 0.5 mM EDTA, washed in complete medium, and then reseeded with 10 µM ROCK inhibitor Y-27693 (ApexBio) overnight, followed by replacement with complete media. WA01 cells are derived from the inner cell mass (ICM) of the blastocyst stage of embryonic development and retain their capacity to differentiate into cells of different lineages[23,24]. Gene expression analysis indicated expression of markers associated with primed pluripotent cells, which includes Sox2, Oct4 and Nanog (Fig S1A) but not markers associated with naïve pluripotency [25]. We confirmed expression of Oct4 and Nanog protein by Western blot and immunofluorescence microscopy (Fig. S1 B, C).

Glioma stem cells (GSC) were grown as previously described [26,27]. Melanoma cells (A101) were maintained as an adherent culture in DMEM media (Cytiva, SH30022.02) supplemented with glutamine analog SG-200 (Cytiva, SH30590.01), 1% sodium pyruvate (Corning, 25-000-CI), and 1% penicillin/streptomycin (Cytiva, SV30010). All cell cultures were maintained at 37 °C and 5% CO_2_ in a humidified atmosphere.

Cells were treated with the following small molecule inhibitors, MEK1/2 inhibitors trametinib (ApexBio, A3018) and PD0325901 (ApexBio, A3013), ERK inhibitor SCH772984 (ApexBio, B5866), EZH2 inhibitor GSK343 (ApexBio, A3449) or the c-Myc:MAX dimerization inhibitor 10058-F4 (ApexBio, A1169).

### RNA extraction and cDNA synthesis

RNA was extracted from cells using RNeasy Mini Kit (Qiagen, 74106) and DNase-digestion was performed during RNA extraction using RNase-free DNase Set (Qiagen, 79254). cDNA was synthesis was primed with random hexamers and oligo dT at final concentration of 1 µM and 5 µM respectively, heated to 65°C for 5 minutes and then snap-cooled. The RNA samples were then reverse transcribed (Protoscript II, New England Biolabs, M0368X) in the prescribed buffer with 0.1 M DTT, 500 mM dNTPs, and RNase inhibitor (Promega, N2511) at 25°C for 5 minutes, 42°C for 60 minutes and 65°C for 20 minutes. Following this, the cDNA samples were treated with RNase H (New England Biolabs, M0297L) and incubated at 37°C for 30 minutes and 65°C for 20 minutes. Quantitative PCR was performed using SYBR Select (ThermoFisher, 4472908) using CFX real-time PCR system (Biorad) and CFX Maestro software was used for quantification. All PCR amplicons were sequenced at least once to ensure accuracy of product obtained. Primers used for amplification as listed in Supplementary Table 1.

### RNA sequencing and data analysis

RNA sequencing was performed by the UAB Genomics Core Lab using 500 ng of RNA as input. Sequencing libraries were made using the NEBNext Ultra II Directional RNA-seq kit (NEB #E7760L) as per the manufacturer’s instructions and using the NEBNext Poly(A) mRNA Magnetic Isolation Module (NEB #E7490). The resulting libraries were analyzed on the BioAnalyzer 2100 (Agilent) and quantified by qPCR (Roche). Sequencing was performed on the NovaSeq 6000 (Illumina) with paired-end 100 bp chemistry as per standard protocols. Fastqs were processed using the nf-core/rnaseq (v3.14.0) pipeline [28] with the --gcBias flag, aligning to the GRCh38.p13 reference, and using the Gencode v32 transcript annotations. Counts were transformed to fragments per kilobase of transcript per million mapped reads (FPKM) for plotting.

### Cell Lysis and Immunoblots

After removing media from cells in a 6-well plate, 1.5 mL of ice-cold PBS was added, and the cells were scraped into a 1.5 mL tube. Cells were collected by centrifugation at 400 × g for 4 min, PBS was removed, and cells were placed on ice for lysis or stored at −80 °C. Cell pellets were lysed in 10 mM Tris-Cl (pH 8.0), 150 mM sodium chloride, 1% Triton X-100, 1 mM EDTA, with Complete Protease Inhibitor (Fisher, A32963) and phosphatase inhibitors (ApexBio, K1015). Samples were incubated on ice for 20 min, then centrifuged for 20 min at 13,000 × g to remove insoluble material. Lowry protein estimation was conducted to quantify the concentrations of cell lysates with DC™ Protein Assay Reagent S (Bio Rad, 500-0115) and DC Protein Assay Reagent A (Bio Rad, 5000113) according to the manufacturer’s protocol. Absorbance readings at 750 nm using a BioTek Synergy 2 plate reader were used to calculate mg/mL protein values. 50 μg of protein were made up in 1 × NuPage LDS sample buffer (Invitrogen, NP0007) and Invitrogen Novex 10 × Bolt Sample Reducing Agent (Thermo Fisher Scientific, B0009), incubated at 95°C for 7 min before loading equal protein amounts onto Mini-PROTEAN TGX Stain-Free Gels 4–20% Tris-Glycine polyacrylamide gels (Bio Rad, 4568096). Gels were run in 1 x TGS buffer for 60 min at 110 volts and transferred to Trans-Blot® Turbo™ Mini Nitrocellulose membranes (Bio Rad, 1704158) for 5 min using the Trans-Blot® Turbo™ RTA Transfer Kit, Nitrocellulose System (Bio Rad, 170-4270), and Trans-Blot Turbo Transfer Buffer (Bio Rad, 10026938). Membranes were blocked in 5% StartingBlock™ (TBS) Blocking Buffer (Fisher, 37542) with orbital shaking for 30 min at room temperature (RT). Primary antibodies were incubated with blots overnight at 4°C with orbital shaking, followed by antibody removal and washing in 10 mL of TBS-T three times for 5 min at RT with orbital shaking. Primary antibodies were detected by the addition of species-specific horseradish-peroxidase conjugated-secondary antibody in 5% nonfat dry milk, incubated with orbital shaking overnight at 4°C followed by washing as for primary antibodies. After removing the last wash buffer, chemiluminescent visualization (SuperSignal™ West Pico PLUS Chemiluminescent Substrate, Thermo Fisher Scientific, 34578) was used. Membranes were visualized on a BioRad ChemiDoc MP. Membranes were then stripped three times in a low-pH stripping buffer (25 mM Glycine, 1% SDS, pH 2.3) for 15 min on an orbital shaker and then washed in 10 mL TBS-T for 5 min on the orbital shaker before the new primary antibody was added.

### Chromatin Immunoprecipitation (ChIP)

Adherent cells in 15 cm plates were washed with PBS then fixed in 1% formaldehyde in PBS for 10 minutes at room temperature, and then treated with 125 mM glycine for 2 minutes to inactivate the formaldehyde. The solution was aspirated, and cells were immediately scraped into 15 mL of ice-cold PBS and spun at 500 x g for 5 min; the PBS was removed and cells were frozen at −80°C. Cells were then lysed for 10 min on ice in 500 mL of 50 mM Tris-Cl pH 8, 10 mM EDTA, 0.5% SDS, complete protease (Fisher, A32963) and phosphatase inhibitors (ApexBio, K1015). Nuclei were disrupted and chromatin solubilized by sonication in a BioRuptor in 1.5 mL tubes until fragments were between 100 and 300 base pairs. The size of fragmented DNA was assessed by running the purified DNA on a 1% agarose gel and visualizing using BioRad GelDoc. Solubilized chromatin was obtained by centrifuging the sonicated lysate at 16,000 x g for 15 min at 16°C and discarding the pellet. The chromatin was quantified using Nanodrop. 10 mg of chromatin were nutated overnight with antibody in 1 mL IP buffer (16.7 mM Tris-Cl pH 8.1, 1.2 mM EDTA, 167 mM NaCl, 1 % Triton X-100) at 4°C. For precipitation, 20 µL of magnetic beads were washed once in IP buffer then added to samples, nutated at 4°C for 24 hours. Chromatin bound to magnetic beads was isolated using a magnetic rack and washed with 1 mL of low-salt buffer (20mM Tris-Cl pH8.0, 2 mM EDTA, 150 mM NaCl, 0.1% SDS, 1% Triton X-100), 1 mL of high-salt buffer (20 mM Tris-Cl pH8.0, 2 mM EDTA, 500 mM NaCl, 0.1% SDS, 1% Triton X-100), 1 mL of LiCl wash (10mM Tris-Cl pH8.0, 1 mM EDTA, 250 mM LiCl, 1% deoxycholate, 1% NP40) and then 1 mL of TE (10 mM Tris pH 8, 2 mM EDTA) for 1 minute each. The TE was removed and the immunoprecipitates were eluted from the beads by incubating in 120 µL of elution buffer (100 mM NaHCO3 and 1% SDS) for 30 minutes at RT, mixing gently every 10 minutes. The supernatant was collected and crosslinks were reversed with 5 mM NaCl at 65°C for 24 hours. The protein and RNA in the samples were digested for 1 h using 7 µL of 1 M Tris pH 6.5, 3 µL of 500 mM EDTA, 3 µL of proteinase K (20 mg/mL), 0.5 µL of RNase A (ThermoFisher, EN5031) and then purified using phenol:chloroform: isoamyl alcohol extraction followed by ethanol precipitation with 0.5 mg glycogen. Input chromatin was purified alongside the immunoprecipitates. Following ethanol precipitation of the purified chromatin, samples were analyzed by quantitative PCR with SYBR Select (ThermoFisher, 4472908) on a CFX real-time PCR system (Bio Rad) and CFX Maestro software was used for quantification. PCR reaction for the *TERT* promoter included 0.5 µL of 7-deaza GTP (Roche, 14020724) per 10 µL reaction. Primers used for amplification as listed in Supplementary Table 1.

### Immunofluorescence microscopy

Imaging was carried out on a Nikon Eclipse Ti inverted confocal microscope equipped with a Nikon 4-line illuminator (405 nm, 488 nm, 561 nm, 640 nm lasers), an X-Cite TURBO 120 LED light source, a Nikon 60x Plan Apo (1.3 NA) oil-immersion objective lens, Nikon 40x Plan Fluor oil-immersion objective (1.30 NA), Nikon 20x PlanApo objective lens (0.750 NA) and a Nikon C2 confocal camera. The microscope was operated using the Nikon Elements software. hESC were collected by EDTA-dissociation and washed with PBS. The cells were resuspended in 1% BSA in PBS and centrifuged onto Superfrost plus slides (Fisher). They were stained simultaneously with mouse anti-nanog antibody at a dilution of 1:250 and rabbit anti-Oct4 antibody at a dilution of 1:500 in 1% BSA in PBS overnight at 4^0^C. The cells were then incubated with two secondary antibodies, anti-rabbit IgG conjugated with DyLight594 and anti-mouse IgG conjugated with DyLight488 at a concentration of 1:500 in 1%BSA in PBS in room temperature for two hours. The slides were imaged as previously described in [26]. Antibodies used are listed in Supplementary Table 2.

### Quantitative Telomerase Repeated Amplification (Q-TRAP) Assay

hESC were collected through EDTA-dissociation, centrifuged at 300 × g for 4 minutes at 4^0^C to pellet. The pellet was washed with PBS. Cells were counted on a Countess (Thermo Fisher) and 500,000 cells per condition were analyzed for telomerase activity using the TRAPeze RT Telomerase Detection Kit (Millipore Sigma. S7710) following manufacturer’s instructions. The cells were lysed in 200 µL of CHAPS lysis buffer containing 200 units/mL of RNase inhibitor (Promega, N2511) on ice for 30 minutes and then centrifuged at 12000 × g for 20 minutes at 4^0^C to collect the supernatant. Telomerase activity was then assessed on 5,000 cell equivalents (2 μl of lysate). In each reaction, the telomerase enzyme catalyzed the addition of telomeric repeats to the 3’ end of the oligonucleotide substrate. Following this the extended products are amplified using the *Taq* DNA polymerase (New England BioLabs, M0273S), and TS and fluorescein labelled Amplifluor primers in the prescribed 5X TRAPeze RT reaction mix. The fluorometric signal resulting from the incorporation of the primers into the PCR products during the reaction was analyzed using CFX Opus 96 real-time PCR system (Bio-Rad) and CFX Maestro software. The standard curve for quantification of telomerase activity in individual samples was obtained by serially diluting the control template TSR8 which is an oligonucleotide with identical sequence to the TS primers and 8 telomeric repeats. The control template was diluted in CHAPS lysis buffer to obtain concentrations ranging from 20 amoles/µL to 0.02 amoles/µL and 2 uL of each was used in similar reaction conditions as above obtain the standard curve.

## RESULTS

### MEK/ERK signaling is required for efficient TERT expression in a model of normal human pluripotent stem cells

Efficient *TERT* expression is essential for stem cells to appropriately set telomere lengths for their differentiated progeny. We previously reported that *TERT* expression in fibroblast-derived human iPSC was dependent on MEK1/2 signaling [29]. To elucidate the effect of MEK1/2 signaling on *TERT* expression in human pluripotent stem cells, we treated hESC and human iPSC with potent, cell-permeable small molecule inhibitors of MEK1 and MEK2 kinase (trametinib or PD0325901). We studied the effect of dosage and duration of MEK1/2 inhibition (MEKi) on *TERT* expression, as assessed by quantitative RT-PCR for exon 2. We found that MEKi decreased *TERT* mRNA levels in hESC (Fig. 1A, B, C). Similar repression of *TERT* was also observed at other *TERT* exons (Fig. S2 A, B). These findings indicate that iPSC and hESC share similar reliance on MEK1/2 signaling for efficient *TERT* transcription.

**Figure 1.**
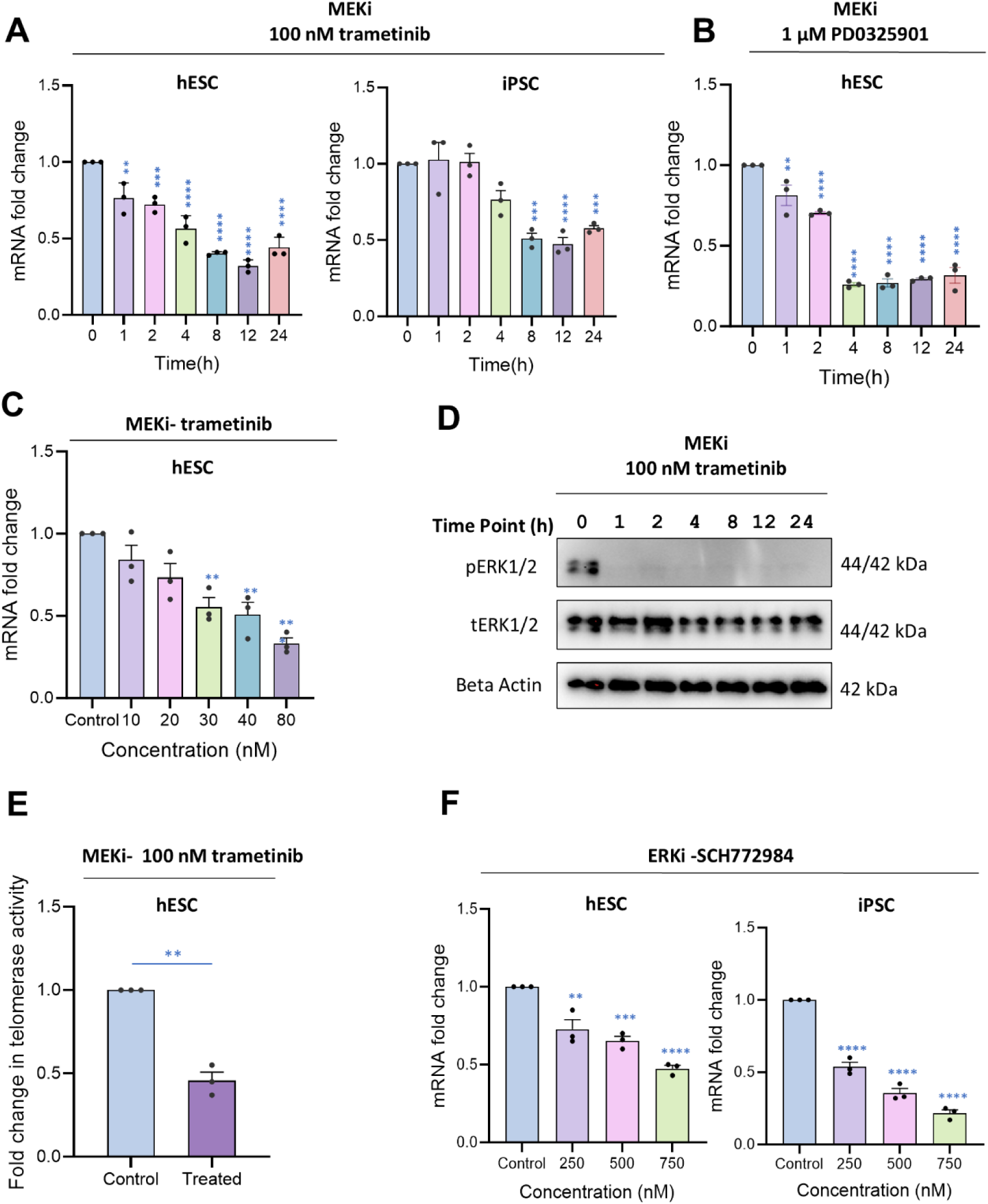
MEK-ERK signaling pathway regulates expression of *TERT* in normal human embryonic stem cells. **A.** *TERT* mRNA levels over the course of 24 hours after treating hESC and iPSC with MEKl/2 inhibitor (MEKi) trametinib. **8.** Expression levels of *TERT* exon 2 mRNA in hESC after treating with an alternate MEKi, PD0325901. **C.** Levels of *TERT* mRNA in hESC cells following treatment with different concentrations of MEKi trametinib for 24 hours. **D.** Western blot showing phospho-ERK (pERK) and total-ERK (tERK) levels in hESC after treatment with trametinib. B-actin was used as a loading control. **E.** Telomerase activity determined by qTRAP assay in cells treated with 100 nM MEKi for 48 hours. **F.** *TERT* mRNA expression in hESC and iPSC after treating with an ERK kinase inhibitor (ERKi) SCH772984. *TERT* expression was assessed by qRT-PCR for exon 2. Graphs depict mean +/− SEM, n=3. *p<0.05, **p<0.01, ***o<0.001 **** o<0.0001. One-wav ANOVA.

Analysis of phosphorylated ERK (pERK) activity via Western blot from cells treated in parallel confirmed loss of ERK activation by MEK1/2, as total ERK levels remained unchanged (Fig. 1D). To determine if reduction in *TERT* mRNA levels following MEK inhibition translates to loss of telomerase activity in cells, we performed quantitative TRAP assay[30]. Telomerase activity in hESC treated with trametinib was significantly reduced 48 hours following treatment (Fig. 1E, Fig. S1D).

To test if the effects of MEK inhibition on *TERT* is mediated via its canonical downstream signaling effectors, ERK1/2, cells were treated with an ERK kinase inhibitor, SCH772984. A dose-dependent decrease in *TERT* expression was observed in hESC and iPSC cells 24 hours after treatment, indicating that efficient *TERT* expression in human pluripotent cells relies on MEK-ERK signaling (Fig. 1F).

### pERK recruitment to the TERT promoter in hESC does not drive MEK1/2 dependent effects

In human embryonic stem cells, pERK2 has been found to bind the active promoters of key pluripotency related genes, in concert with transcription factors such as ELK1, and play an important role in regulating the pluripotent identity of these cells [31]. Given the sensitivity of *TERT* transcription to MEK-ERK inhibition in both stem cells (Fig. 1) and some cancer cells [25,35,36] we posited a potential shared ERK-dependent mechanism controlling the *TERT* promoter in both cancer cells and stem cells. For example, in melanoma cells, pERK is recruited to the *TERT* promoter but is lost upon MEK inhibition [33]. We first examined the effect of MEK1/2 inhibition on pERK recruitment to the *TERT* promoter in melanoma and glioma stem cells (GSC) using chromatin immunoprecipitation (ChIP). Our results confirmed a previous report that MEKi causes loss of pERK from the *TERT* promoter in melanoma cells [36] and extends these observations to GSCs (Fig. 2A). However, contrary to expectations, no significant changes in pERK levels at the *TERT* promoter in hESC and iPSC were observed upon MEK inhibition (Fig. 2B), suggesting that the role of MEK/ERK in regulating *TERT* in stem cells differs from cancer cells. Of note, the level of pERK bound to the *TERT* promoter in GSC and melanoma cells was markedly higher than that observed in stem cells. This disparity further suggests distinct MEK/ERK-dependent mechanisms regulating *TERT* expression in cancer cells compared to human pluripotent stem cells.

**Figure 2.**
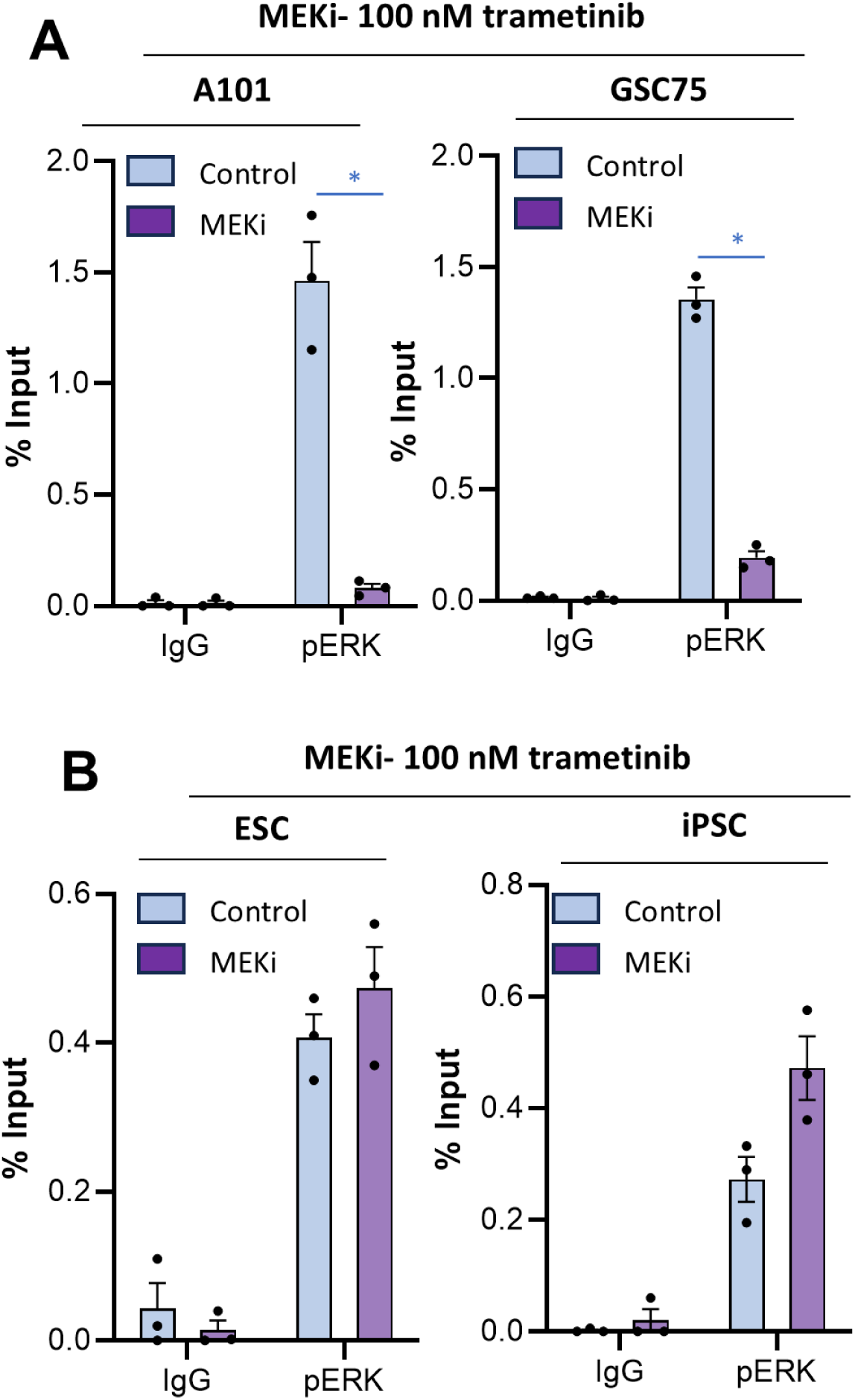
Inhibiting RAS signaling fails to deplete activated ERK (pERK) from the *TERT* promoters in hESC but not in cancer cells. **A.** ChlP-qPCR performed in melanoma cell line AlOl, glioblastoma cell line GSC75 showing pERK enrichment at the *TERT* promoter after treatment with MEKi trametinib for 24 hours. **B.** ChlP-qPCR performed in hESC and iPSC showing ERK levels at *TERT* promoter after treatment with MEKi (trametinib) for 24 hours. The graphs depict % input, +/−SEM, n=3 independent biological replicates. *p<0.05, **p<0.01,

### MEK1/2 activity partly functions to prevent polycomb repression by PRC2 at TERT

Since loss of pERK from the *TERT* promoter did not explain the reduction in *TERT* mRNA expression in stem cells after MEK inhibition, we explored alternative mechanisms for *TERT* repression following MEK inhibition. We first examined levels of H3K27ac at the *TERT* proximal promoter, a mark that is associated with active transcription, and H3K27me3, a histone mark associated with transcriptionally repressed chromatin. After trametinib treatment, we observed a decline in H3K27ac at the *TERT* proximal promoter (Fig. 3A). This was accompanied by increased trimethylation of histone 3 proteins at lysine 27 (H3K27me3). These data suggest that the loss of acetylation allowed for deposition of methyl groups on the same histone proteins. Trimethylation of histone 3 lysine 27 is catalyzed by EZH2, the enzymatic subunit of polycomb repressive complex 2 (PRC2). Thus, our results suggest that one function of MEK1/2 kinase activity in driving *TERT* is to limit the deposition of H3K27me3. To test this idea, we treated hESC with an EZH2 inhibitor (GSK343) at the same time as a MEK inhibitor. In these cells, *TERT* expression could be partially rescued by inhibiting EZH2 (Fig. 3B) indicating that MEK1/2 signaling limits PRC2-mediated repression of the *TERT* promoter.

**Figure 3.**
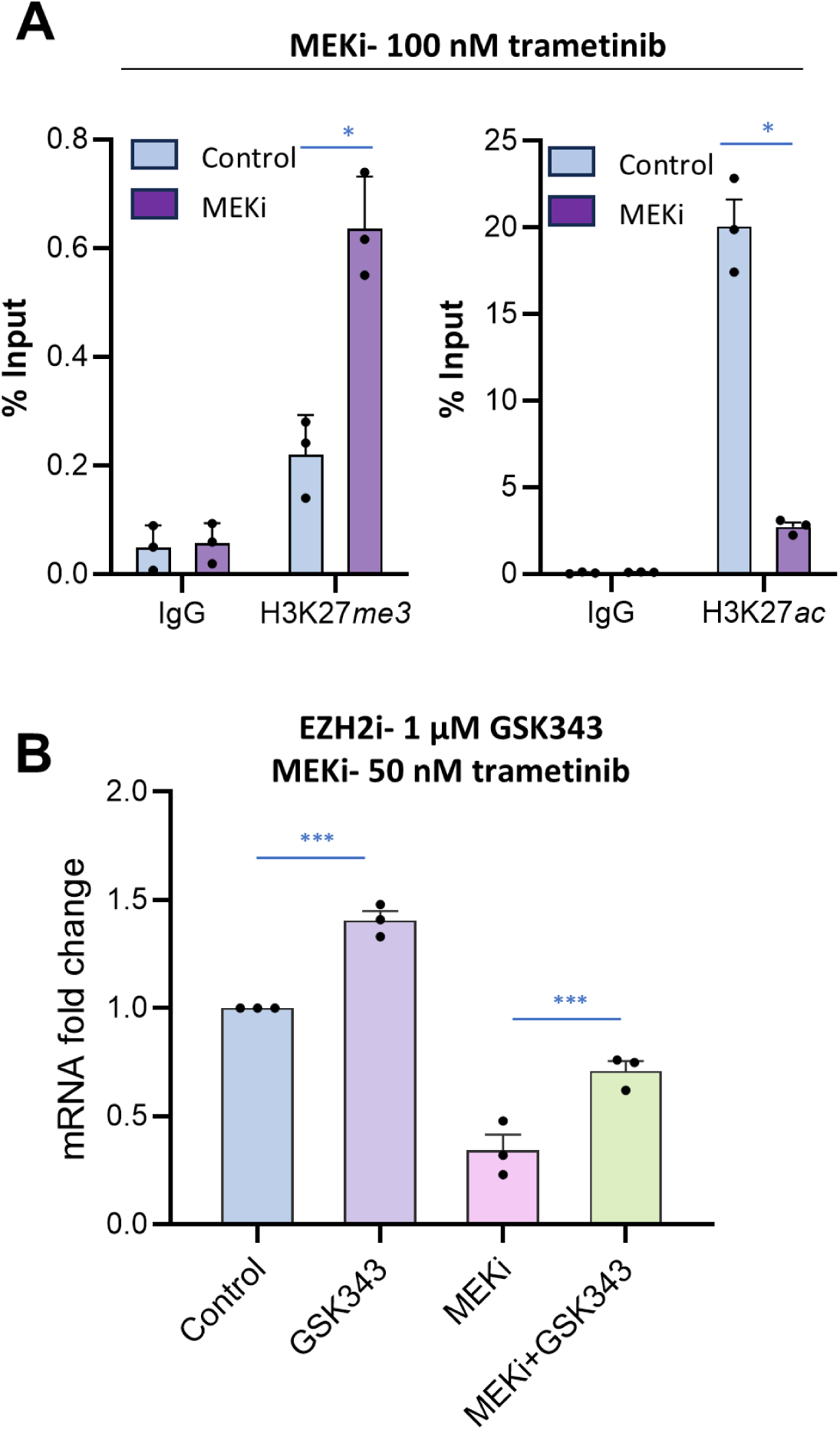
Endogenous RAS signaling promotes histone 3 lysine 27 acetylation at the *TERT* promoter limiting polycomb repressive marks. **A.** ChlP-qPCR for repressive histone mark H3K27me3 and active histone mark H3K27ac at *TERT* promoter in hESC treated with or without 100 nM of MEKi trametinib for 24 hours. The graphs depict % input chromatin pulled down, +/− SEM, n=3 independent biological replicates. **B.** *TERT* mRNA levels in hESC following treatment with 1 µM of EZH2 inhibitor GSK343, SO nM trametinib, or a combination of SO nM trametinib and 1 µM of EZH2 inhibitor GSK343 for 24 hours,. TERT expression was determined by qRT-PCR for exon 2. The graphs depict mRNA fold change, +/− SEM, n=3 independent biological replicates. *p<0.0S, **p<0.01, ***p<0.001 **** p<0.0001 Two-way ANOVA.

### RAS signaling promotes TERT expression in hESC through c-Myc:MAX dimers

As MEK1/2 activity regulates *TERT* transcripts via PRC2 independent of pERK recruitment to the *TERT* promoter, we considered that this control may be mediated via other transcription factors. The human *TERT* promoter contains binding motifs for various transcription factors, including c-Myc, which can regulate *TERT* expression [34–36]. Given that c-Myc expression is also controlled by MEK-ERK signaling in fibroblasts and cancer cells [39,40], we investigated the impact of MEK inhibition on c-Myc expression in hESC. We observed a time-dependent reduction in c-Myc mRNA levels following MEK inhibition, with an 85% decrease noted after 24 hours of trametinib exposure (Fig. 4A). This repression of c-Myc was found to be ERK-dependent, as treatment with the ERK inhibitor, SCH772984, resembled the effects of MEK inhibition (Fig. 4A). These findings suggest that MEK1/2 signaling in hESC may drive *TERT* transcription via c-Myc. To determine if c-Myc is present at the *TERT* promoter, we analyzed publicly available ENCODE ChIP-seq data from H1-ESC [39,40]. Consistent with our hypothesis, we observed c-Myc binding at the endogenous human *TERT* promoter (Fig. S3), indicating that these expression changes in c-Myc may impact c-Myc recruitment specifically at *TERT*.

**Figure 4.**
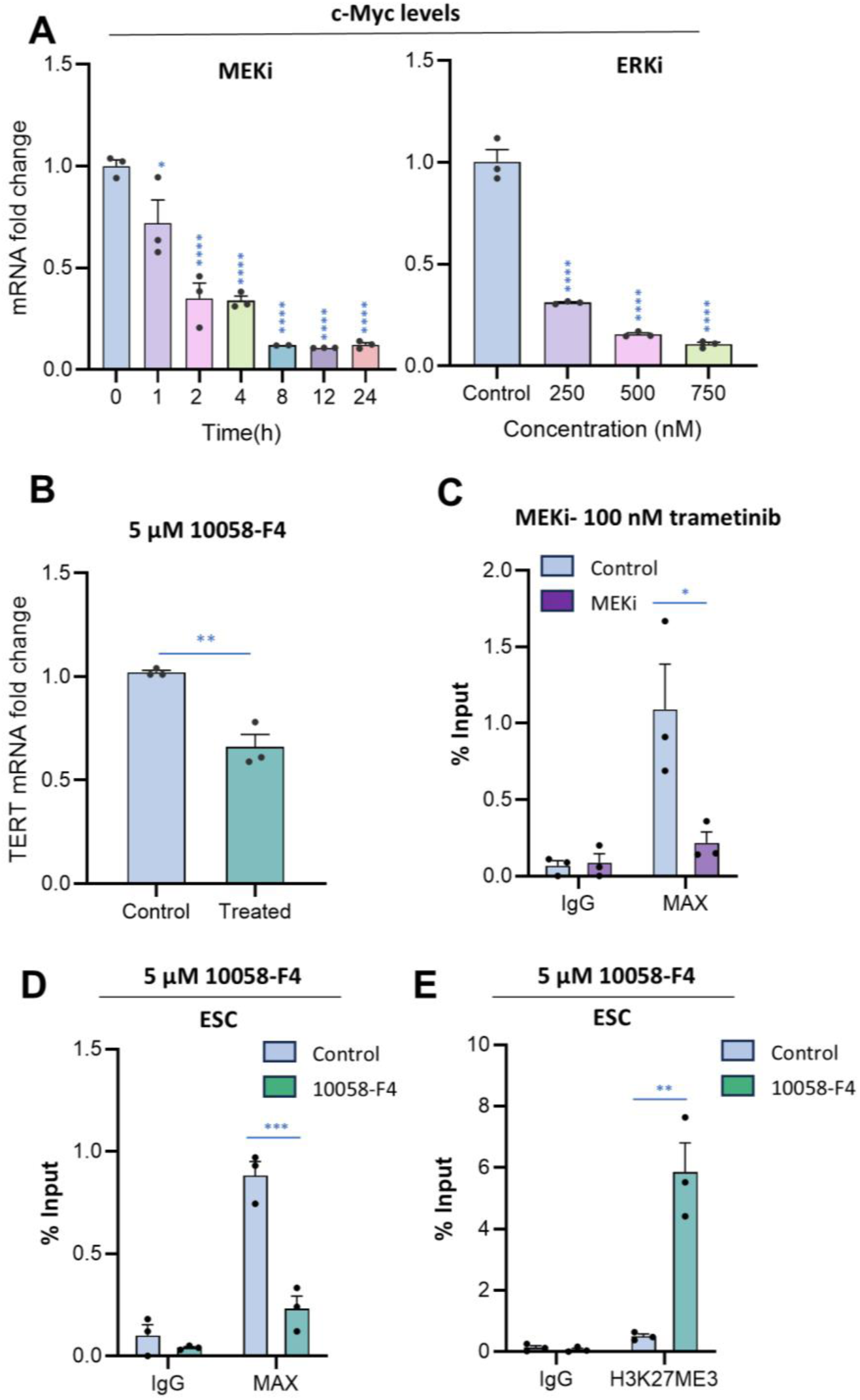
RAS driven *TERT* expression proceeds because c-Myc:MAX dimers limit PRC2/EZH2. **A.** c-Myc mRNA levels in hESC following treatment with MEKi trametinib (100 nM) or ERKi SCH772984 for 24 h. Expression was determined by qRT-PCR. The graphs depict mRNA fold change, +/− SEM, n=3. **B.** *TERT* mRNA levels in hESC treated with c-Myc:MAX dimerization inhibitor 10058-F4 at 5 **µM** for 8 hours. **C.** ChlP-qPCR for MAX in hESC following 24 h treatment with MEKi (trametinib 100 nM). **D.** ChlP-qPCR for MAX in hESC following 8 h treatments with 10058-F4 (5 **µM). E.** ChlP-qPCR for repressive histone mark H3K27me3 at the *TERT* promoter in hESC following treatment with 10058-F4 (5 **µM)** for 12 hours. Graphs in C-E depict % Input chromatin pulled down, +/−SEM, n=3 independent biological replicates. *p<0.05, **p<0.01, ***p<0.001 **** p<0.0001, Two-way ANOVA.

### c-Myc:MAX dimers mediate RAS dependent inhibition of polycomb repression of TERT

c-Myc forms heterodimers with the protein MAX to bind to DNA sequences [41,42]. We therefore hypothesized that the effect of c-Myc on *TERT* transcription may be dependent on this dimerization. Examination of ENCODE ChIP-seq data indicates that MAX does bind to the proximal *TERT* promoter (Fig. S3). To test whether dimerization may regulate binding of MAX to *TERT*, we took advantage of a well characterized, selective small molecule inhibitor of c-Myc:MAX dimerization, 10058-F4 [43–45]. Following an 8-hour exposure to low levels (5 µM) of this inhibitor, a significant reduction in *TERT* expression was observed in hESC (Fig. 4B). This suggested that c-Myc binding at *TERT* promoter in combination with MAX is important for efficient *TERT* expression in hESC cells. To test if c-Myc:MAX heterodimers act in cis at the *TERT* promoter, we performed ChIP experiments with anti-MAX antibody following treatments with MEK inhibitor trametinib or 10058-F4 in hESC. Decreased MAX levels at the *TERT* promoter were evident in both cases (Fig. 4C, D), further substantiating the importance of c-Myc:MAX heterodimer in *TERT* transcription. Consistent with this rapid decrease in *TERT* transcription, ChIP analysis of the *TERT* promoter after treatment with c-Myc:MAX dimerization inhibitor revealed a striking and rapid gain of the repressive histone mark H3K27me3 (Fig. 4E).

## DISCUSSION

In this study, using a model of normal human pluripotent stem cells, we demonstrate that transcription of the endogenous human *TERT* gene is under the control of MEK1/2 and ERK1/2 kinases. Inhibition of MEK1/2 kinase activity was accompanied by elevated levels of the H3K27me3 catalyzed by EZH2, the enzymatic subunit of PRC2. Inhibition of EZH2 activity led to partial rescue of *TERT* transcript levels. MEK inhibition also caused significant decreases in c-Myc expression and loss of MAX from the *TERT* promoter and formation of c-Myc:MAX dimers was essential for efficient *TERT* transcription in hESC (Fig. 5).

**Figure 5.**
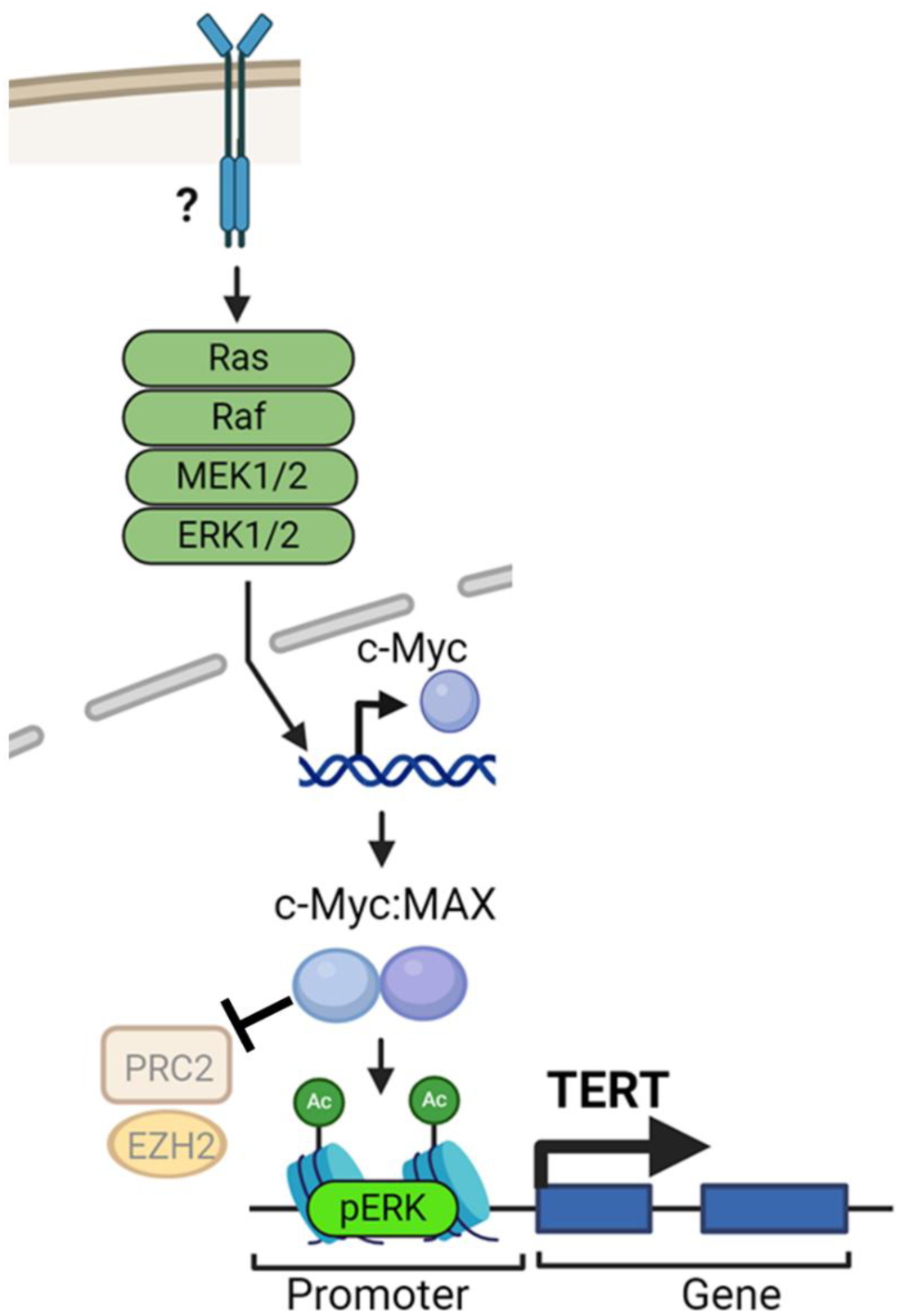
Schematic representation of *TERT* transcriptional regulation MEK-ERK pathway and c-Myc:MAX dimers in hESC. MEKl/2 activation from an unknown upstream signal promotes c-Myc expression, which dimerizes with MAX at *TERT* in hESC to limit PRC2 deposition of H3K27me3. These events coincide with activation of TfRTtranscription.

Studies of exogenous c-Myc or *TERT* promoter constructs suggest that c-Myc has the capacity to drive *TERT* expression in stem cells [34,46]. However, a regulatory role for endogenous c-Myc in normal human pluripotent stem cells at the native human *TERT* locus has not been demonstrated. Our data indicate that c-Myc:MAX dimers are essential for efficient *TERT* transcription. Inhibition of dimerization was coupled with a striking gain of H3K27me3 mark at the *TERT* promoter, revealing a role for c-Myc:MAX dimers at *TERT* in preventing PRC2 mediated repression. PRC2 is an enzyme involved in silencing developmental genes in embryonic stem cells and maintaining pluripotency [47,48]. Future studies will be aimed at deciphering the molecular pathways involved in EZH2 inhibition by c-Myc:MAX.

An interesting outcome from our study is the observation that MAX binding was lost from the *TERT* promoter following either MEK inhibition or c-Myc:MAX dimerization inhibition. *TERT* has been reported to have two canonical E-boxes at the promoter which are potential binding sites for basic helix-loop-helix leucine zipper (bHLH-LZ) family of transcription factors which includes c-Myc:MAX. Additionally, intron 2 of the *TERT* gene consists of a variable number tandem repeat which also consists of dozens of E-boxes. Both c-Myc and MAX have been shown to be enriched at intron 2 in bacterial artificial chromosome reporter construct of the *TERT* gene in a mouse embryonic cell line [49]. Future studies will be aimed at testing the downstream transcription mechanisms regulated by these MAX binding events.

Investigations of phosphorylated ERK have revealed its occupancy at transcriptionally active promoters of numerous genes pivotal for proliferation, survival, and self-renewal [31]. Our ChIP experiments with normal human pluripotent stem cells also revealed pERK occupancy at the *TERT* promoter. Intriguingly, our study identified differences in pERK dynamics following MEK inhibition in pluripotent stem cells compared to cancer cells. Specifically, in cancer cells pERK is lost upon MEKi treatment. However, in hESC, despite global loss of pERK in cells, no change was observed at *TERT*. This finding suggests that even though some cancer cells are also sensitive to MEK inhibitors, it may partly occur through divergent mechanisms from those operating in stem cells. One possible explanation for pERK persistence at stem cell *TERT* promoters in the presence of MEKi or ERKi is that they exhibit different dephosphorylation dynamics of pERK bound to chromatin in cancer cells. Future studies examining expression or activity of nuclear phosphatases targeting chromatin bound ERK may reveal mechanistic differences in stem cells vs cancer cells. One class of phosphatases, the dual specificity phosphatases (DUSP), especially DUSP 1, 2, 4 and 5, which are nuclear-specific phosphatases that target ERK, could be responsible. Studying the expression pattern and/or post-translational modifications of these phosphatases may hold the key to determining differential binding dynamics of chromatin bound pERK at *TERT*.

Several studies have reported cell culture methods to model naïve pre-implantation epiblasts. These media conditions commonly include MEK inhibitors which promote an undifferentiated state [50–53]. Our study raises interesting questions regarding telomere maintenance and the *TERT* transcription under these conditions. Does *TERT* expression rely on MEK1/2 signaling under naïve conditions? If so, given that *TERT* is haploinsufficient, our data suggest that naïve cells cultured for long periods with MEK inhibitors may suffer from telomeric defects. Consistent with this hypothesis, it was recently observed that MEK inhibition in naïve human ESC promoted genomic instability [54]. That study, as well as our data, suggest that alternatives to MEK inhibitors for promoting an undifferentiated state of naïve stem cells may prove beneficial.

For this study, we chose a human embryonic stem cell line derived from the blastocyst stage as a model because (i) this is the stage at which *TERT* is activated during development, and (ii) normal human developmental tissue cannot be readily examined. Consequently, these cell types represent the best available model. Observations that were tested in iPSC (derived from a different donor and sex) were similar to the hESC, indicating they are likely to be a general feature of *TERT* transcriptional regulation in human pluripotent stem cells.

The concentrations we have used in this study for trametinib and the Myc:MAX dimer inhibitor are low and similar to those used for many prior studies. These compounds are extensively characterized and validated by previous work, lending strong confidence that results we observe are from on-target effects.

In conclusion, our study provides a picture of the interplay of epigenetic and transcriptional regulators coordinated by MEK-ERK signaling to maintain *TERT* transcription in human pluripotent stem cells to ensure their long-term proliferative capacity.

