## Supplemental Information for "MEK1/2 kinases cooperate with c-Myc:MAX to prevent polycomb repression of *TERT* in human pluripotent stem cells"

Kotian et al 2024
Supplementary Materials
Figs. S1 - S3
Tables S1, S2

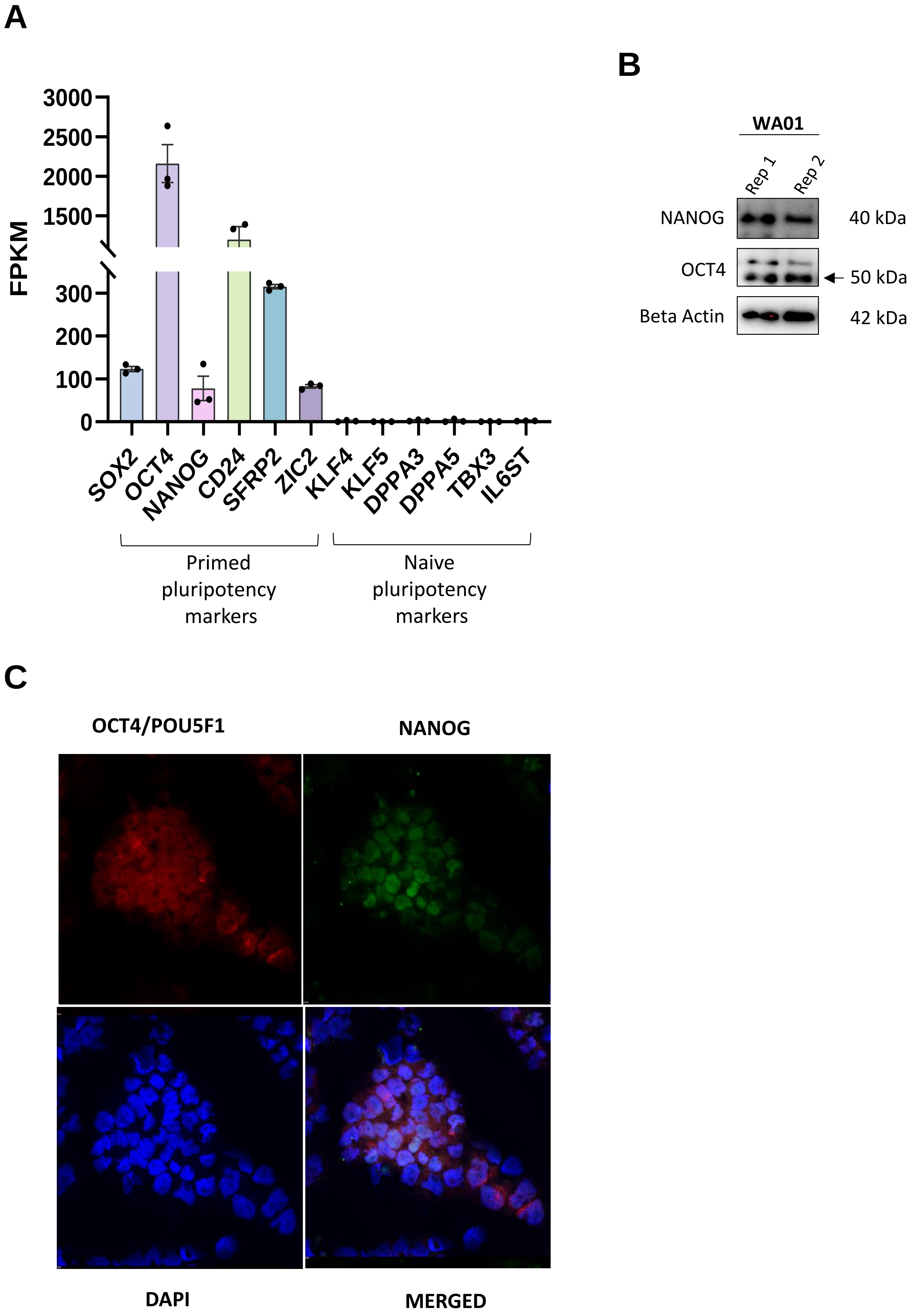

**Figure S1. WA01 human embryonic stem cells display markers of primed pluripotency. A.** RNA-seq data for WA01 cells analyzed for different markers of pluripotency. Data are fragments per kilobase of transcript per million mapped reads (FPKM) +/- SEM. **B.** Western blot showing NANOG and OCT4 levels in hESC. ß-actin is used as a loading control. **C.** Immunofluorescence images of hESC cells stained with anti-NANOG, anti-OCT4 antibody, and DAPI.

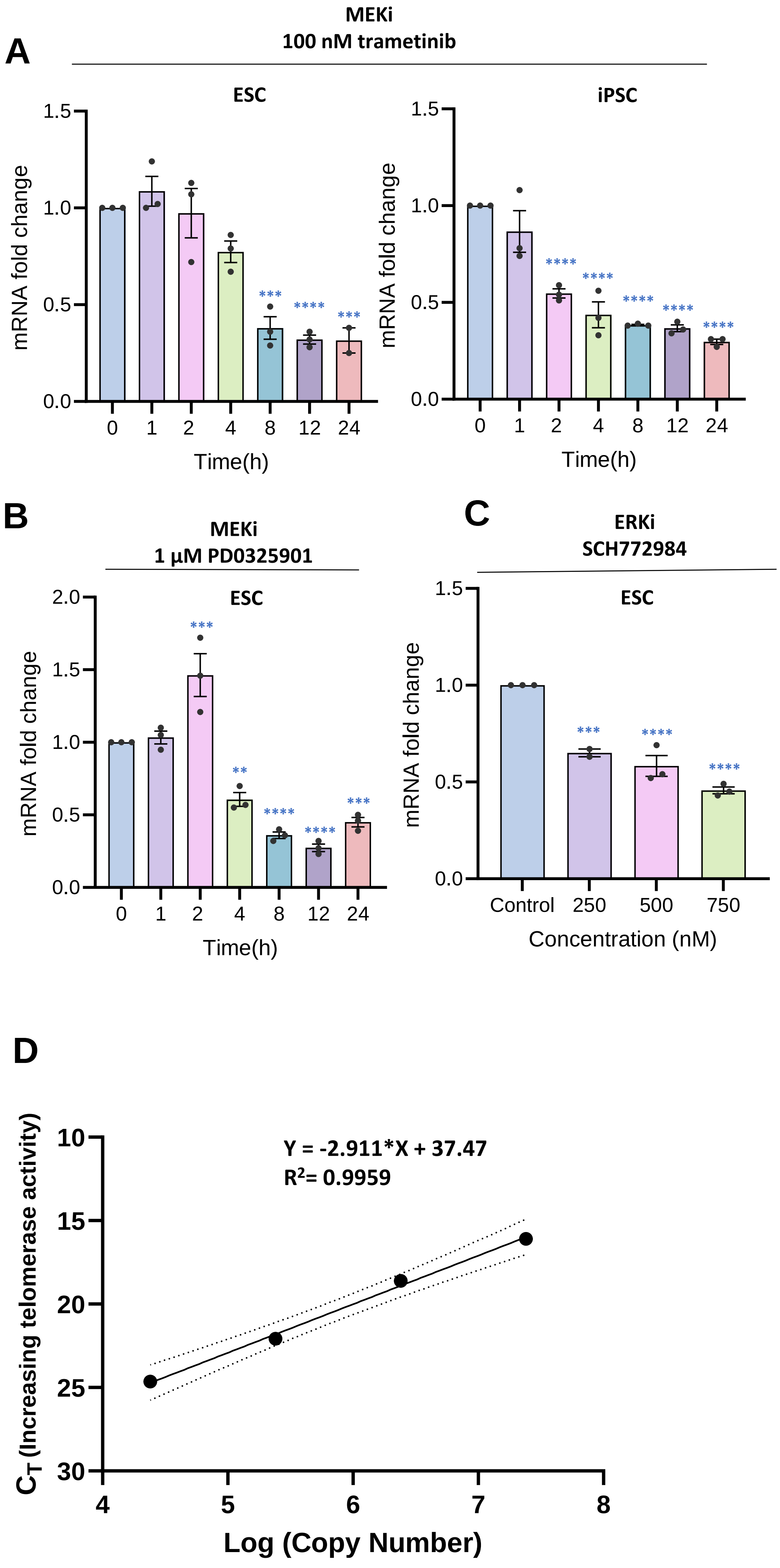

**Figure S2. *TERT* exon 14 levels following inhibition of MEK-ERK signaling . A.** *TERT* mRNA levels assessed by qRT-PCR for exon 14 over the course of 24 hours after treating hESC and iPSC with MEK1/2 inhibitor (MEKi) trametinib. **B.** Expression levels of *TERT* mRNA in hESC after treating with an alternate MEKi, PD0325901 determined by qRT-PCR for exon 14. **C.** *TERT* exon 14 mRNA expression in hESC after treating with an ERK kinase inhibitor (ERKi) SCH772984. Graphs depict means with +/- SEM for n=3. *p<0.05, **p<0.01, ***p<0.001 **** p<0.0001 Ordinary one-way ANOVA. **D.** Logarithmic plot of the TSR8 standard curve for quantification of telomerase activity in Fig 1E. Graph depicts the CT values corresponding fluorometric signal obtained on the amplification of different dilutions of TSR8 template.

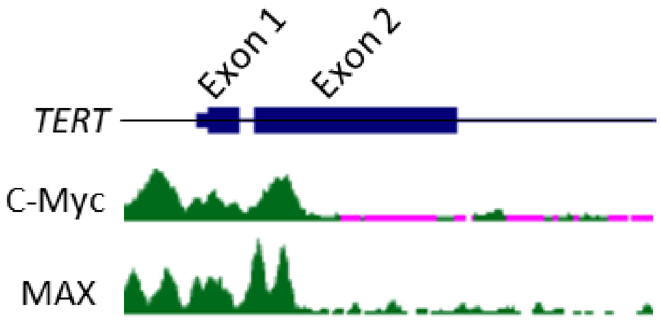

**Figure S3**. **Myc and MAX recruit to *TERT* in WA01 hESC.** Figure displays occupancy of c-Myc and MAX proteins at the *TERT* locus as determined by ChIPseq. Data for c-Myc (GSM935509) were from the Snyder lab at Stanford; for MAX (GSM1010898) were from Myers Lab, Hudson Alpha Institute.

| | | |
|---|---|---|
| TERT exon 2 | Forward | CTACTCCTCAGGCGACAAGG |
| | Reverse | TGGAACCCAGAAAGATGGTC |
| TERT exon 14 | Forward | CATTTCATCAGCAAGTTTGGAAG |
| | Reverse | TTTCAGGATGGAGTAGCAGAGG |
| OCT4 | Forward | GGTCCGAGTGTGGTTCTGTA |
| | Reverse | GCAGCCTCAAAATCCTCTCG |
| NANOG | Forward | ACCCCAGCCTTTACTCTTCC |
| | Reverse | CTGGATGTTCTGGGTCTGGT |
| TERT promoter | Forward | GTCCTGCCCCTTCACCTT |
| | Reverse | AGCGCTGCCTGAAACTCG |
| GAPDH | Forward | CTGCACCACCAACTGCTTAG |
| | Reverse | GTCTTCTGGGTGGCAGTGAT |
| c-Myc | Forward | TACCCTCTCAACGACAGCAG |
| | Reverse | CGTCGAGGAGAGCAGAGAAT |

**Supplementary table 1.** List of primers

| | | |
|---|---|---|
| **Antibodies for western blot** | | |
| p-ERK | Cell Signaling Technology | 9101L |
| t-ERK | Cell Signaling Technology | 9102S |
| Actin | Santa Cruz Technology | sc-47778 |
| Oct4 | ProteinTech | 11263-1-AP |
| NANOG | ProteinTech | 14295-1-AP |
| **Antibodies for ChIP** | | |
| IgG | Cell Signaling Technology | 2729S |
| H3K27me3 | Active Motif | 91403 |
| H3K27ac | Cell Signaling Technology | 8173S |
| p-ERK | Cell Signaling Technology | 9101L |
| MAX | GeneTex | GTX113492 |
| **Antibodies for IF** | | |
| Rabbit Oct4/POU5F1 anti-Oct4 | ProteinTech | 11263-1-AP |
| Mouse anti-nanog | AbCam | ab173368 |
| Goat Anti-rabbit IgG (DyLight594) | GeneTex | GTX213110-05 |
| Goat Anti-mouse IgG (DyLight488) | GeneTex | GTX213111-04 |

**Supplementary table 2.** List of antibodies
